# Novel biomaterial-based synovial fluid analysis reveals protective microRNA signatures in a mouse model of acute synovitis-driven osteoarthritis

**DOI:** 10.1101/2025.09.11.675532

**Authors:** Kei Takahata, Kohei Arakawa, Takashi Yasuda, Saaya Enomoto, Takuya Katashima, Takamasa Sakai, Takanori Kokubun

**Affiliations:** Graduate School of Health, Medicine, and Welfare, Saitama Prefectural University, Saitama, Japan; Laboratory of Animal Regeneration Systemology, Department of Life Sciences, School of Agriculture, Meiji University, Kanagawa, Japan; Faculty of Advanced Life Sciences, Hokkaido University, Hokkaido, Japan; Saitama Prefecture General Rehabilitation Center, Saitama, Japan; Graduate School of Engineering, The University of Tokyo, Tokyo, Japan; Department of Physical Therapy, School of Health and Social Services, Saitama Prefectural University, Koshigaya, Saitama, Japan

**Keywords:** Osteoarthritis, Synovial fluid, Synovitis, Mechanical stress, miRNA, biomaterials

## Abstract

Acute synovitis and continuous mechanical stress are critical factors of Post-traumatic Osteoarthritis (PTOA); however, the effect on cartilage degeneration and its underlying mechanisms remains unclear. Here, we established a novel biomaterial-based method for synovial fluid analysis in mice to uncover the mechanisms underlying the onset of PTOA, focusing on synovitis and mechanical stress. Twelve-week-old C57BL/6J males were divided to the ACL-Transection (ACL-T), ACL-rupture (ACL-R), and Intact groups. We performed joint instability test and histological analysis for cartilage degeneration and synovitis at 2, 6, and 10 weeks. Real time PCR was performed on articular cartilage at 2 weeks and synovium at 2, 6, and 10 weeks. Tetra-slime was injected into the knee joint, and solidified slime containing synovial fluid molecules was analyzed in digital PCR at 2 weeks. Acute synovitis and cartilage degeneration were induced in the ACL-T group at 2 weeks, although no difference was observed in joint instability between both two ACL injury models. Real time PCR showed significant increase of Mmp-3, Tnf-α, Ifn-γ, and inos in synovium of the ACL-T group. During this tissue interaction, although there were no significant differences, miR145-5p and miR149-5p in synovial fluid were also upregulated in the ACL-T group compared to the ACL-R group. Unlike the early stage, no histological or biological differences were observed between groups at 6 and 10 weeks. In conclusion, acute synovitis caused secondary cartilage degeneration via MMP-3 and TNF-α in the synovium during the early stage and may involve M1 macrophage activation. Whereas it was suggested that synovial fluid miRNA might be produced to suppress the secondary cartilage degeneration. Furthermore, mechanical stress was the dominant factor in the late stage of OA, regardless of the initial synovitis condition.

## Introduction

Post-traumatic Osteoarthritis (PTOA) is characterized by whole-joint disease following joint trauma such as meniscal and ligament injuries. PTOA induces cartilage degeneration, synovitis, and subchondral bone sclerosis, and these structural changes influence each other through synovial fluid and progress irreversibly. ^1^ Acute inflammation with tissue injuries and continuous mechanical stress due to joint instability are known as the PTOA onset factors. Previous studies reported that inflammation following joint trauma, especially synovitis, might be the initiator of OA, resulting in cartilage degeneration through the innate immune system. ^2–4^ However, it is challenging to understand both effects of acute synovitis and chronic mechanical stress separately in patients. To elucidate the PTOA onset mechanism in terms of inflammation and mechanical stress, an assessment of synovial fluid, which is a key mediator within the knee joint, is essential in the animal models.

Synovial fluid is an ultrafiltrate of blood plasma and is primarily composed of hyaluronan and lubricin, which play a significant role as a joint space lubricant and a nutrient source for intra-articular tissues. ^5^ Previous studies have reported that Interleukin (IL)-6, IL-8, and Tumor Necrosis Factor (TNF)-α in synovial fluid are correlated with OA severity ^6^, and Matrix Metalloproteinase (MMP)-3 could be a biomarker for detecting early-stage OA. ^7^ Furthermore, circulating microRNAs (miRNAs) in the synovial fluid have been highlighted as a potential biomarker to reflect the intra-articular environment. ^8^ miRNAs are short, non-coding RNAs that are typically 18-25 nucleotides in length, which are used to investigate various diseases since they are stable compared to other RNAs degraded by ribonucleases.^9^ It has been reported that miR-23a-3p, miR-24-3p, miR-27b-3p, miR-29c-3p, miR-34a-5p, and miR-186-5p in the synovial fluid were expressed in the late-stage OA compared to the early-stage. ^10^ In addition, miR-210 increased significantly in the OA synovial fluid than that in healthy individuals. ^11^ Many clinical studies have investigated synovial fluid changes to reveal the onset mechanism of OA in humans ^12^, which has yet to be done due to the variability of symptoms and disease progression in the early stages of OA. ^13, 14^

Preclinical experiments using animal OA models are crucial for understanding human pathologies, and many mouse models have been used. ^15, 16^ Generally, joint lavage was performed to collect synovial fluid in murine knee joints as follows: 25-50 µL of saline was initially injected into the knee joint, and then the synovial fluid was drawn out. ^17–19^ However, it is challenging to collect a sufficient amount of synovial fluid and present relevant data due to the tiny knee joints in mice. ^20^ Recently, various types of biomaterials have been developed and applied in drug delivery, tissue engineering, and medical diagnostics. ^21–23^ Tetra-slime is composed of reversible boronic ester bonds between phenylboronic acid and diol groups, and its stiffness can be changed by adjusting the precursor concentration and pH. ^24^ Moreover, it can also be dissolved by sorbitol as sugars attach to phenylboronic acid and cleave ester bonds. ^25^ Based on these material properties, we hypothesized that harvesting and dissolving the solidified Tetra-Slime, which was injected into the knee joint, would allow for the efficient collection of synovial fluid. The purpose of this study was to elucidate each effect of acute synovitis and continuous mechanical stress on PTOA progression using surgical Anterior Cruciate Ligament Transection (ACL-T) and Non-Invasive ACL rupture (ACL-R) mouse models. Moreover, we proposed a novel method to analyze synovial fluid miRNAs in murine knee joints using Tetra-slime to reveal these tissue interactions.

## MATERIALS AND METHODS

### Animals and Experimental Design

This study was approved by the Animal Research Committee of Saitama Prefectural University (approval number: 2020-6). The animals were handled according to relevant legislation and institutional guidelines for humane animal treatment. Seventy-two 12-week-old C57BL/6 male mice were randomized into the Invasive ACL-T group (n = 36) and Non-Invasive ACL-R group (n = 36). The contralateral knee joint in the ACL-R group was used as the control group. Six mice in each group were sacrificed at 2, 6, and 10 weeks, and the target tissues were collected for each analysis (Fig. 1). All mice were housed in plastic cages under a 12-hour light/dark cycle. The mice were allowed unrestricted movement within the cage and had free access to food and water.

**Fig. 1.**
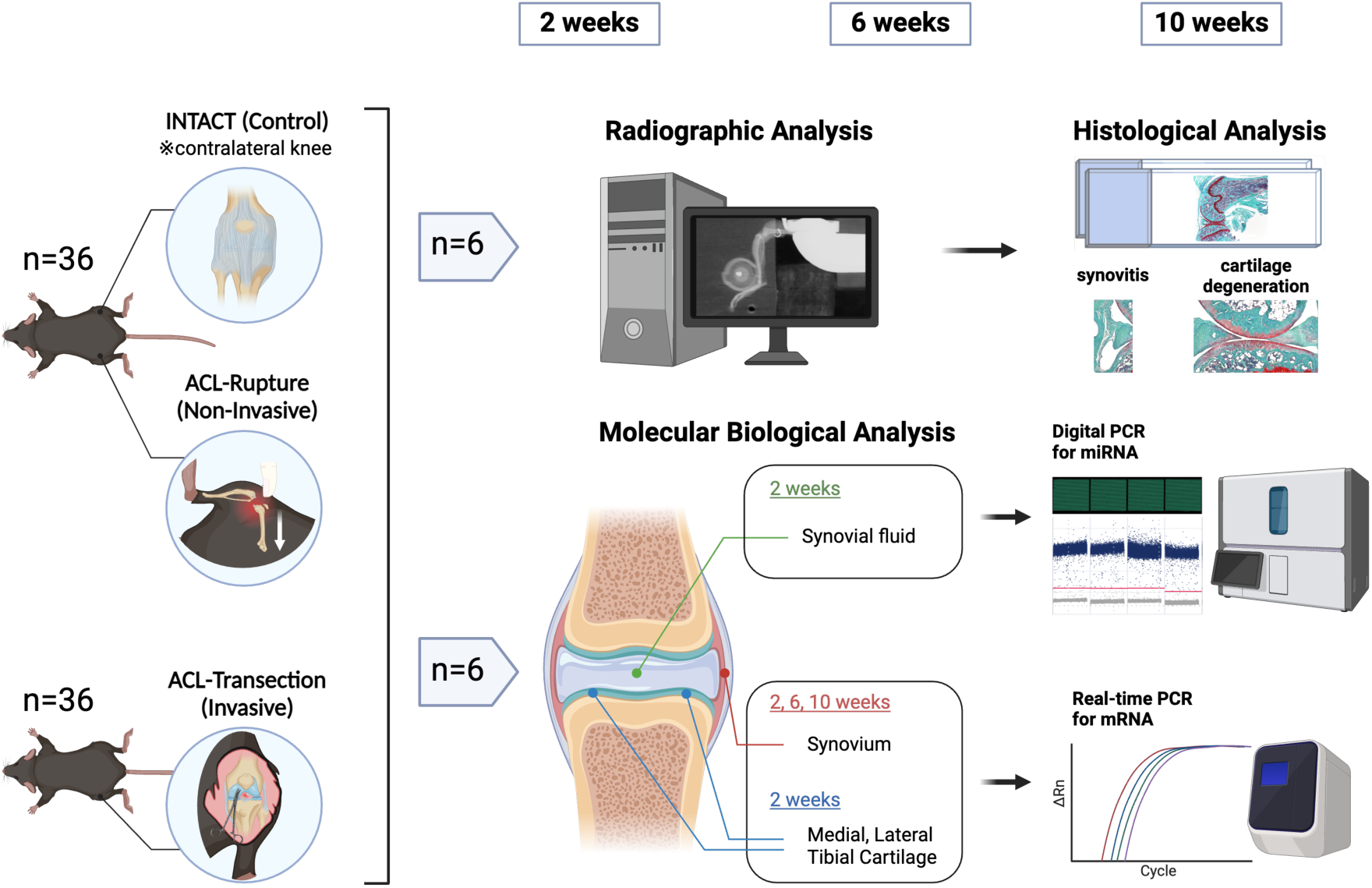
Experimental design. We made the Invasive ACL-T, Non-Invasive ACL-R, and Control groups. Contralateral knee in the ACL-R model was used as Control group. At 2, 6, and 10 weeks, anterior drawer test and histological analysis for synovitis and cartilage denegation in the medial and lateral compartments were performed. Articular cartilage at 2 weeks and synovium at 2, 6, and 10 weeks were harvested, and mRNAs were analyzed in real time PCR. In addition, synovial fluid was collected at 2 weeks and miRNAs level were assessed.

### Creating ACL-T and ACL-R models

All procedures were performed on the left knee joint of each mouse under a combination of anesthetics (medetomidine, 0.375 mg/kg; midazolam, 2.0 mg/kg; and butorphanol, 2.5 mg/kg). The medial joint capsule was cut to expose the intra-articular joint. ACL was transected with scissors to avoid causing injuries to other tissues. After checking the anterior tibial translation manually, the joint capsule and skin incision were closed with 5-0 nylon threads. The ACL-R model was created based on our previous study.^26^ Briefly, the knee joint was fixed at 90 degrees using surgical tape on a stand, then the force along the long axis of the femur was applied by the thumb’s tip slowly and carefully. Applied force was stopped quickly after hearing a distinct popping sound, which indicates ACL rupture; thereafter, anterior tibial translation was confirmed manually as well.

### Ex vivo radiographic analysis for knee joint instability

After dissection at each time point, knee joints were collected (each group, n = 6) and an anterior drawer test was performed to evaluate joint instability (Fig. 2A). Based on our previous study ^16, 26^, the proximal tibia was pulled forward with a constant force spring (0.05 kgf; Sanko Spring Co., Ltd., Fukuoka, JPN), and X-ray images were taken at this position. In the acquired image, the tibial tuberosity and the anterior end of the femoral condyle were set as landmarks, then the anterior tibial displacement was measured by the linear distance from the perpendicular line from the anterior end of the femoral condyle to the tibial tuberosity using Image J (National Institutes of Health, Bethesda, MD, USA).

**Fig. 2.**
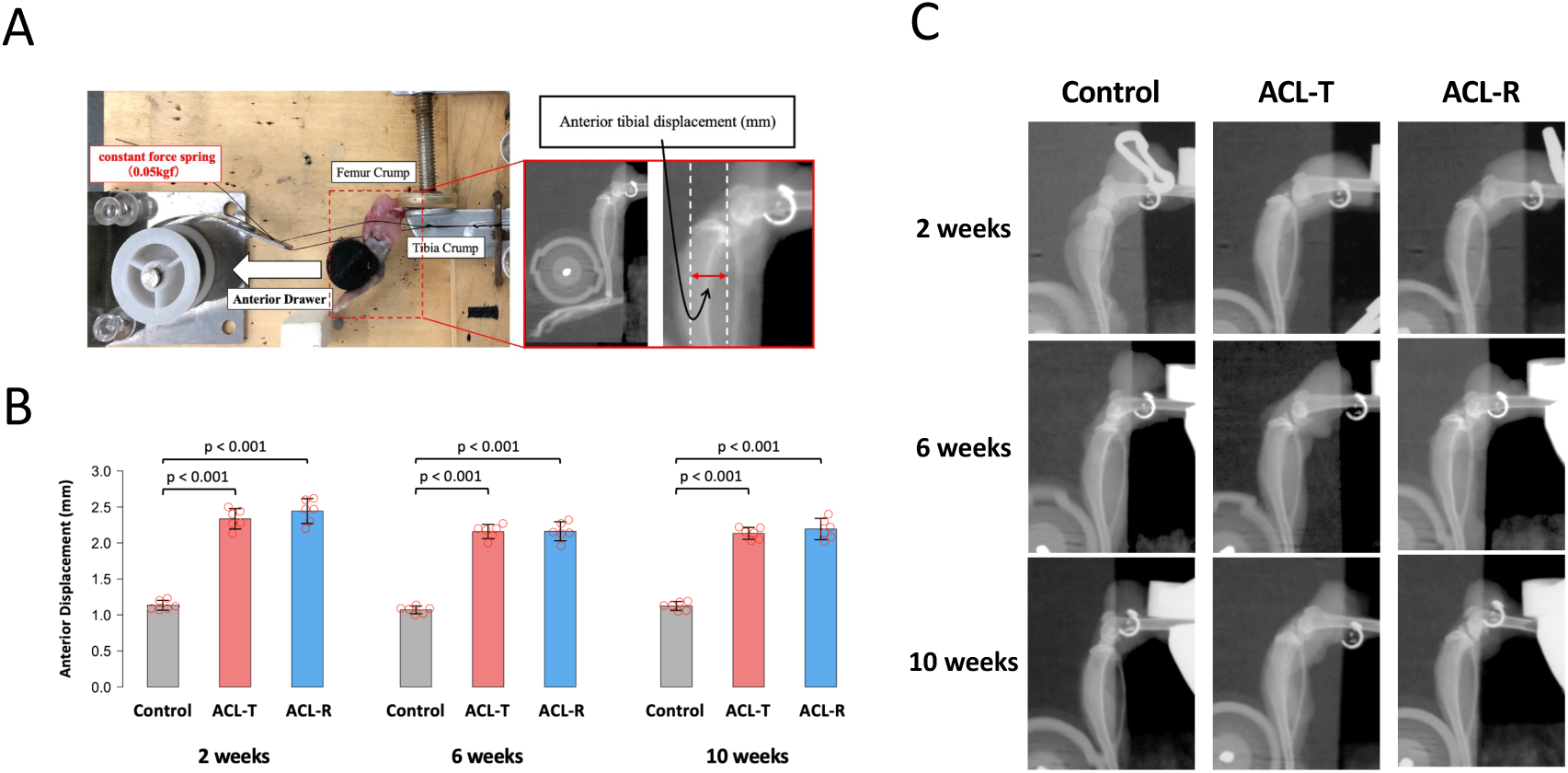
Evaluation of the joint instability *ex vivo* using soft X-ray and original anterior drawer device. At 2, 6, and 10 weeks, the amount of anterior tibial displacement was significantly increased in the ACL-T and ACL-R groups compared to that in the Control group. However, there was no significant difference between both ACL injury models. Data are presented as the mean ± 95% CI.

### Histological analysis for cartilage degeneration and synovitis

Knee joints were fixed using 4% paraformaldehyde for 1 day, decalcified in 10% ethylenediaminetetraacetic acid for 2 weeks, dehydrated in 70% and 100% ethanol and xylene, and embedded in paraffin blocks after performing radiographic analysis. The samples were cut in the sagittal plane (thickness, 7 µm) using a microtome (ROM-360; Yamato Kohki Industrial Co., Ltd., Saitama, JPN). Then, safranin-O/fast green staining was performed to assess the articular cartilage degeneration with the Osteoarthritis Research Society International (OARSI) histopathological grading system ^27^ and synovitis score ^28^ in the medial and lateral compartments. These scores were assessed by two independent observers blinded to all other sample information. We evaluated the contact area not covered by the meniscus on the tibial plateau and the synovium located inside the infrapatellar pad. The mean of the observer’s scores was used as a representative value.

### Real-time PCR for articular cartilage and synovium

Articular cartilage on the medial and lateral tibial plateau at 2 weeks and synovium at 2, 6, and 10 weeks were collected and stored in a deep freezer until analysis (each group, n = 6). Based on the manufacturer’s instructions, total RNA was extracted from these tissues using the ISOGEN (NIPPON GENE CO., LTD., Tokyo, JPN). cDNA synthesis was performed from the total RNA by Thermal Cycler Palm-Cycler (Corbett Research, SYD, AUS) with High-Capacity cDNA Reverse Transcription kit (Applied Biosystems, CA, USA). Subsequently, real-time PCR was conducted to assess the expression of following genes using the StepOne-Plus system (Applied Biosystems, Waltham, MA, USA); *Mmp-3*(Mm00440295), Tissue inhibitor of metalloproteinase-1 (*Timp-1:* Mm01341361), Transforming growth factor-β1 (*Tgf-β1:* Mm01178820), *Tnf-α* (Mm00443258), *Il-1β (*Mm00434228), *Il-6* (Mm00446190), *Il-10* (Mm01288386), Interferon-γ (*Ifn-γ* :Mm01168134), inducible nitric oxide synthase (*inos* :Mm00440502_m1), and Toll like receptor-4 (*Tlr4* :Mm00445273_m1). We used Glyceraldehyde-3-phosphate dehydrogenase (*Gapdh* :Mm99999915) as a housekeeping gene, and relative expression of genes was detected by the 2-△△Ct method. ^29^

### Digital PCR for synovial fluid miRNA using Tetra-Slime

In this study, we selected the Tetra-slime to collect the murine synovial fluid. 4-arm PEG BFA and 4-arm PEG Gluconic (XIAMEN SINOPEG BIOTECH CO., LTD., Fujian, CHN) were resolved with 50 mM phosphate-buffered saline (pH 3.0) to adjust 0.2 g/ml concentration respectively. We injected both solutions by 25 µL into the knee joint for live mice, then kept the needle for a while to avoid leaking the solution (Fig. 7A). After 3 minutes, mice were sacrificed and the quadriceps and patellar were removed to expose the intra-articular joints, then solidified Tetra-Slime was collected. We added 100 µL of sorbitol solution with a 0.5 g/ml concentration to the collected Tetra-Slime and stirred slowly to dissolve and separate from the surrounding intra-articular tissue, then obtained the pure solution with synovial fluid biomolecules released. Based on the manufacturer’s instructions, miRNA was extracted from the synovial fluid solution using the miRNeasy Micro Kit (QIAGEN, LTD. Tokyo, JPN). cDNA synthesis was performed by QIAamplifier 96 (QIAGEN, LTD. Tokyo, JPN) with miRCURY LNA RT Kit (QIAGEN, LTD. Tokyo, JPN), then digital PCR (dPCR) was conducted using the QIAcuity One (QIAGEN, LTD. Tokyo, JPN) and target genes were *hsa-miR-145-5p* (YP00204483) and *hsa-miR-149-5p* (YP00204321). We normalized the target gene with the housekeeping gene, 5S rRNA (YP00203906), for each sample, and then calculated the relative expression level by standardizing with the Control group. Using the UniSp2 (YP00203950) and UniSp6 (YP00203954) as a spike-in control, the effect of Tetra-slime on miRNA expression, the efficiency of collecting synovial fluid RNA from murine knee joint, and the effect of pH 3.0 on miRNA expression were also explored.

### Statistical analysis

Statistical analysis was performed using RStudio version 1.4.1106. For the results of anterior tibial displacement by the anterior drawer test, relative expression of genes in synovium and cartilage by real-time PCR, and miRNA data measured using dPCR, one-way analysis was performed initially, then the Tukey-Kramer test was used as a post-hoc test. We utilized the value of △Ct for the real-time PCR data and the relative expression value for the dPCR data. Whereas the Kruskal-Wallis test was used to compare the OARSI and synovitis scores, and the Steel-Dwass method was used for the subsequent multiple comparisons.

Parametric data are expressed as the means ± 95% confidence intervals (CI); in contrast, the non-parametric data are expressed as the medians ± interquartile ranges. Statistical significance was set at *p* < 0.05. CIs were shown in the Supplementary data.

## Results

### ACL-T and ACL-R groups showed similar joint instability

The anterior displacement in the ACL-T and ACL-R groups was significantly higher than that in the Control groups at 2 weeks (ACL-T vs. Control, p < 0.001, ACL-R vs. Control, p < 0.001), 6 weeks (ACL-T vs. Control, p < 0.001, ACL-R vs. Control, p < 0.001), and 10 weeks (ACL-T vs. Control, p < 0.001, ACL-R vs. Control, p < 0.001). However, there was no significant difference between the ACL-T and ACL-R groups at all time points. These results suggested that the effect of mechanical stress on OA progression was identical in both ACL-deficient models.

### ACL-T group induced acute synovitis in the early stage

For the medial compartment, severe synovitis with increased cellularity, enlargement of the synovial lining cell layer, and inflammatory infiltration was observed in the ACL-T group (Fig. 3A). Moderate synovitis with cell multiplication and cell layer enlargement was confirmed in the ACL-R group. Compared with the Control group, synovitis score was significantly higher in the ACL-T group (p = 0.009) and ACL-R group (p = 0.012) (Fig 3B). Interestingly, ACL-T group increased score significantly compared with the ACL-R group (p = 0.032). At 6 weeks, moderate synovitis characterized by some enlargement and increased cellularity developed in both ACL-deficient groups, and the synovitis score in the ACL-T group (p = 0.01) and ACL-R group (p = 0.01) was significantly higher than in the Control group. At 10 weeks, similar moderate to severe synovitis was observed in both ACL-deficient groups, and the synovitis score in the ACL-T group (p = 0.009) and ACL-R group (p = 0.01) was significantly elevated compared to the Control group.

**Fig. 3.**
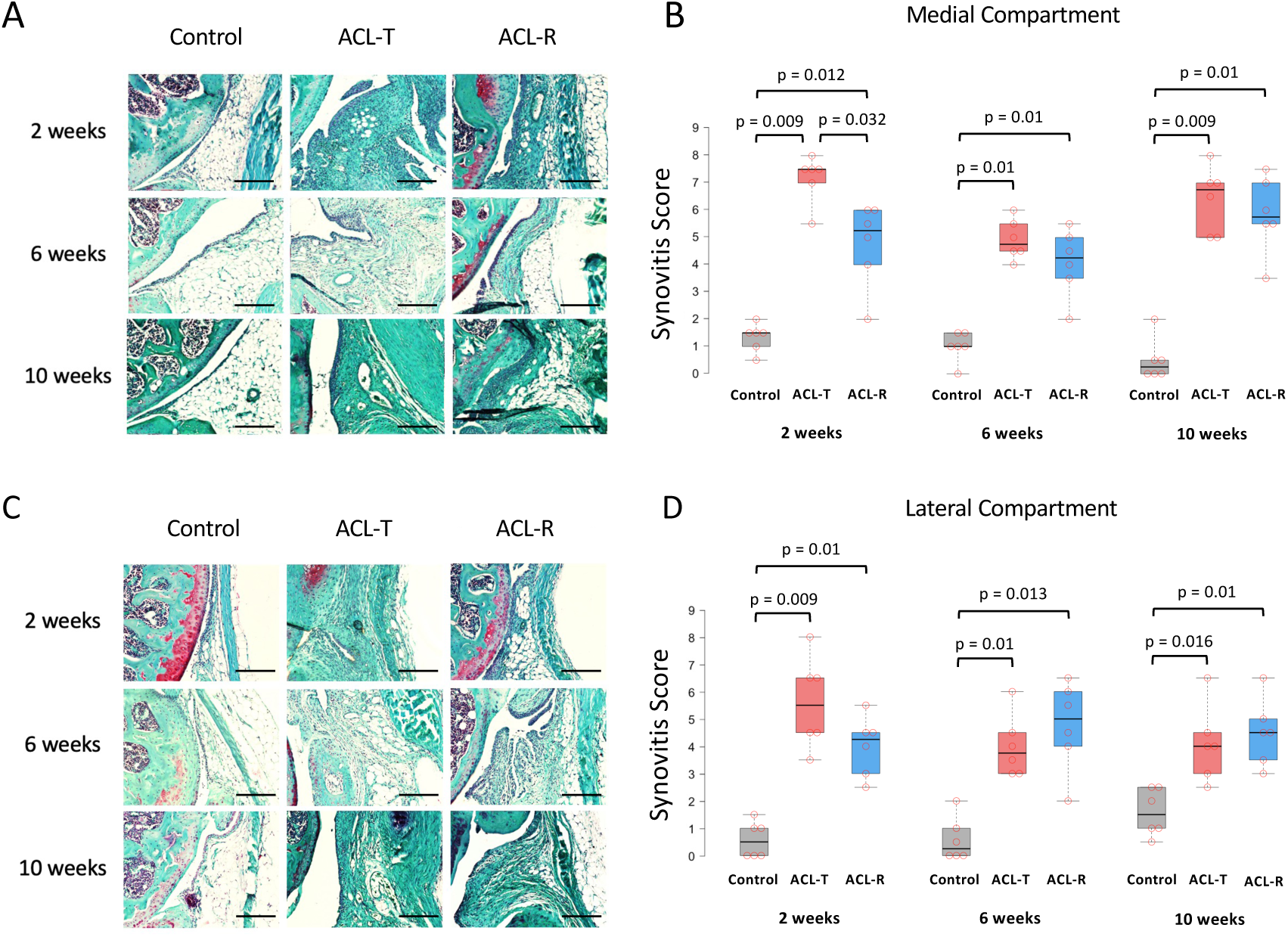
Histological analysis of synovitis using safranin-O/fast green staining and synovitis score. (A-B) In the medial compartment, the synovitis score in the ACL-T and ACL-R groups was significantly higher than that in the Control group at 2 weeks. Moreover, the score in the ACL-T group was also elevated significantly compared to the ACL-R group. At 6 and 10 weeks, both ACL injury groups significantly increased the synovitis score compared with the Control group. (C-D) In the lateral compartment, the ACL-T and ACL-R groups exhibited a significant increase of synovitis score compared to the Control group at all timepoints. Data are presented as the median ± interquartile range. Black scale bar, 100 µm.

For the lateral compartment, moderate and mild synovitis occurred in both ACL-deficient groups. Synovitis score was significantly higher in the ACL-T group (p = 0.009) and ACL-R group (p = 0.01) compared with the Control group (Fig. 3C-D). At 6 and 10 weeks, moderate synovitis was observed in the ACL-T and ACL-R groups, and both synovitis scores increased significantly compared with the Control group (6 weeks: ACL-T vs. Control, p = 0.01; ACL-R vs. Control, p = 0.013, 10 weeks: ACL-T vs. Control, p = 0.016; ACL-R vs. Control, p = 0.01).

### Secondary cartilage degeneration was induced by acute synovitis in the ACL-T group

To examine the tissue interaction between early synovitis and cartilage degeneration, histological analysis was performed. For the medial compartment, some fibrillation of the superficial cartilage was observed in the ACL-T and ACL-R groups at 2 weeks (Fig. 4A).

**Fig. 4.**
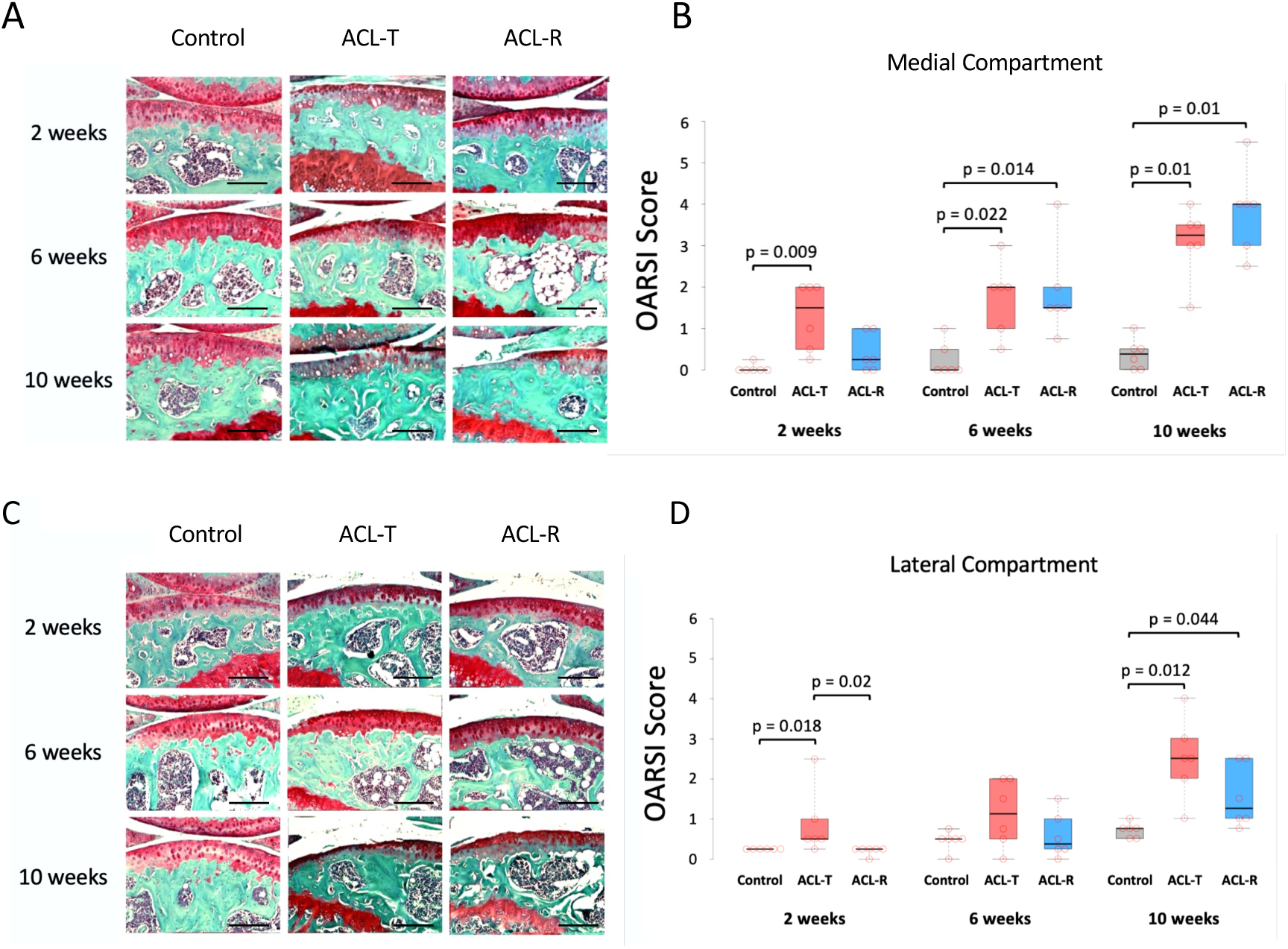
Histological analysis of cartilage degeneration using safranin-O/fast green staining and OARSI score. (A-B) In the medial compartment, the OARSI score in the ACL-T group was significantly higher than that in the Control group at 2 weeks. At 6 and 10 weeks, both ACL injury groups significantly increased the OARSI score compared with the Control group. (C-D) In the lateral compartment, the OARSI score in the ACL-T group increased significantly compared the Control and ACL-R groups at 2 weeks. At 10 weeks, both ACL injury groups exhibited a significant increase of OARSI score compared to the Control group. Data are presented as the median ± interquartile range. Black scale bar, 100 µm.

Additionally, small clefts and loss of cartilage were confirmed only in the ACL-T group. The OARSI score of the ACL-T group was significantly higher than that of the Control group (p = 0.009) (Fig. 4B). At 6 weeks, cartilage fibrillation and mild to moderate vertical cracks into the calcified layer developed in both ACL-deficient groups. Compared with the Control group, the OARSI score was significantly elevated in the ACL-T group (p = 0.022) and ACL-R group (p = 0.014). At 10 weeks, moderate to severe cartilage degeneration with loss of the superficial layer was observed in both ACL-deficient groups. Compared with the Control group, the OARSI score significantly increased in the ACL-T group (p = 0.01) and ACL-R group (p = 0.01).

For the lateral compartment, cartilage degeneration with loss of safranin staining, small fibrillation, and vertical clefts was confirmed in the only ACL-T group at 2 weeks (Fig. 4C). The OARSI score in the ACL-T group was significantly higher compared with that in the Control group (p = 0.018) and ACL-R group (p = 0.02) (Fig. 4D). At 6 weeks, small fibrillation and clefts in the superficial cartilage layer occurred in both ACL-deficient groups; however, there was no significant difference in the OARSI score between all groups. At 10 weeks, mild cartilage degeneration with small fibrillation and vertical clefts in the superficial cartilage layer was observed in both ACL-deficient groups. Compared with the Control group, the OARSI score was significantly higher in the ACL-T group (p = 0.012) and ACL-R group (p = 0.044). In summary of the histological analysis, the ACL-T group induced acute synovitis and cartilage degeneration in the first 2 weeks compared with the ACL-R group, although no difference in the joint instability. Cartilage degeneration in both ACL-deficient models progressed in the late stage, as well as synovitis.

### ACL-R group induced catabolic factors and inflammation cytokine expression in the articular cartilage

To explore the molecular mechanism between acute synovitis and cartilage degeneration, we performed real-time PCR for the articular cartilage at 2 weeks because cartilage degeneration occurred in the whole area moderately after 6 weeks. In the medial tibial cartilage compared with the Control group, ACL-T group significantly increased the expression of *Mmp-3* (p < 0.001), *Timp-1* (p = 0.009), and *Tnf-α* (p = 0.008) (Fig. 5A-C). ACL-R group also significantly exhibited an increase of *Mmp-3* (p < 0.001), *Timp-1* (p < 0.001), and *Tnf-α* (p < 0.001) compared with the Control group. Moreover, *Tnf-α* in the ACL-R group was significantly higher than in the ACL-T group (p = 0.005) (Fig. 5C).

**Fig. 5.**
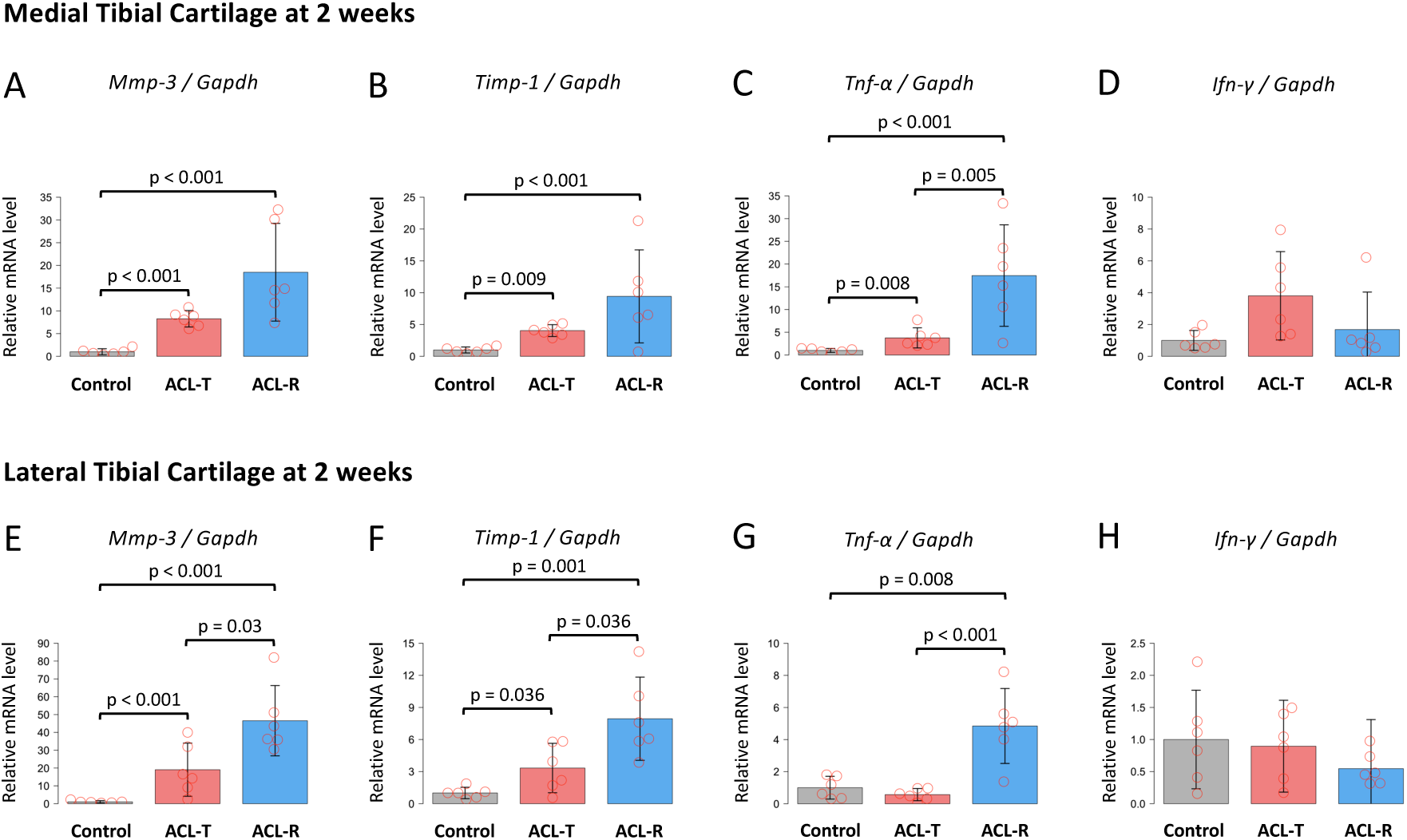
Real time PCR analysis for medial and lateral tibial articular cartilage at 2 weeks. (A-D) In the medial cartilage, *Mmp-3*, *Timp-1*, and *Tnf-α* expression in the ACL-T and ACL-R groups were significantly increased compared to that in the Control group. In addition, the ACL-R group exhibited a significant increase of *TNF-α* compared to the ACL-T group. (E-H) In the lateral cartilage, the expression of *Mmp-3*, *Timp-1*, and *Tnf-α* in both ACL injury groups were significantly upregulated compared to that in the Control group. *Mmp-3* and *Timp-1* were also significantly increased in the ACL-R group compared to the ACL-T group. Data are presented as the mean ± 95% CI.

In the lateral tibial cartilage, ACL-T group significantly exhibited an increase of *Mmp-3* (p < 0.001) and *Timp-1* (p = 0.036) compared with the Control group (Fig. 5E-F). *Mmp-3*, *Timp-1*, and *Tnf-α* in the ACL-R group were significantly upregulated compared with the Control group (*Mmp-3*; p < 0.001, *Timp-1*; p < 0.001, *Tnf-α*; p = 0.008) and ACL-T group (*Mmp-3*; p = 0.03, *Timp-1*; p = 0.036, *Tnf-α*; p < 0.001) (Fig. 5E-G). However, no significant differences were observed in the *Ifn-γ* between all groups for the medial and lateral tibial cartilage (Fig. 5D, H). *Tgf-β1*, *Il-1β*, *Il-6*, and *Il-10* were exhibited in the Supplementary Figure 1.

### ACL-T group increased the expression of catabolic factors, inflammatory cytokines, and M1 macrophage markers in the synovium

At 2 weeks, *Mmp-3* expression was significantly higher in the ACL-T group (p < 0.001) and ACL-R group (p < 0.001) than the Control group (Fig. 6A). Furthermore, the ACL-T group also exhibited an increase of *Mmp-3* expression significantly compared with the ACL-R group (p = 0.048). *Timp-1* was elevated significantly in the ACL-T (p = 0.002) and ACL-R group (p < 0.001) compared with the Control group (Fig. 6B). The ACL-T group significantly increased *Tnf-α*, *Ifn-γ*, and *inos* compared with the Control (*Tnf-α*; p = 0.001, *Ifn-γ*; p = 0.021, *inos*; p = 0.017) and ACL-R group (*Tnf-α*; p = 0.004, *Ifn-γ*; p = 0.012, *inos*; p = 0.009) (Fig. 6C-E). However, no significant difference was observed in the *Tlr4* expression among groups (Fig. 6F). The results for *Tgf-β1*, *Il-1β*, *Il-6,* and *Il-10* at 2 weeks, all results at 6 weeks, and 10 weeks were described in the Supplementary Figures 2-4.

**Fig. 6.**
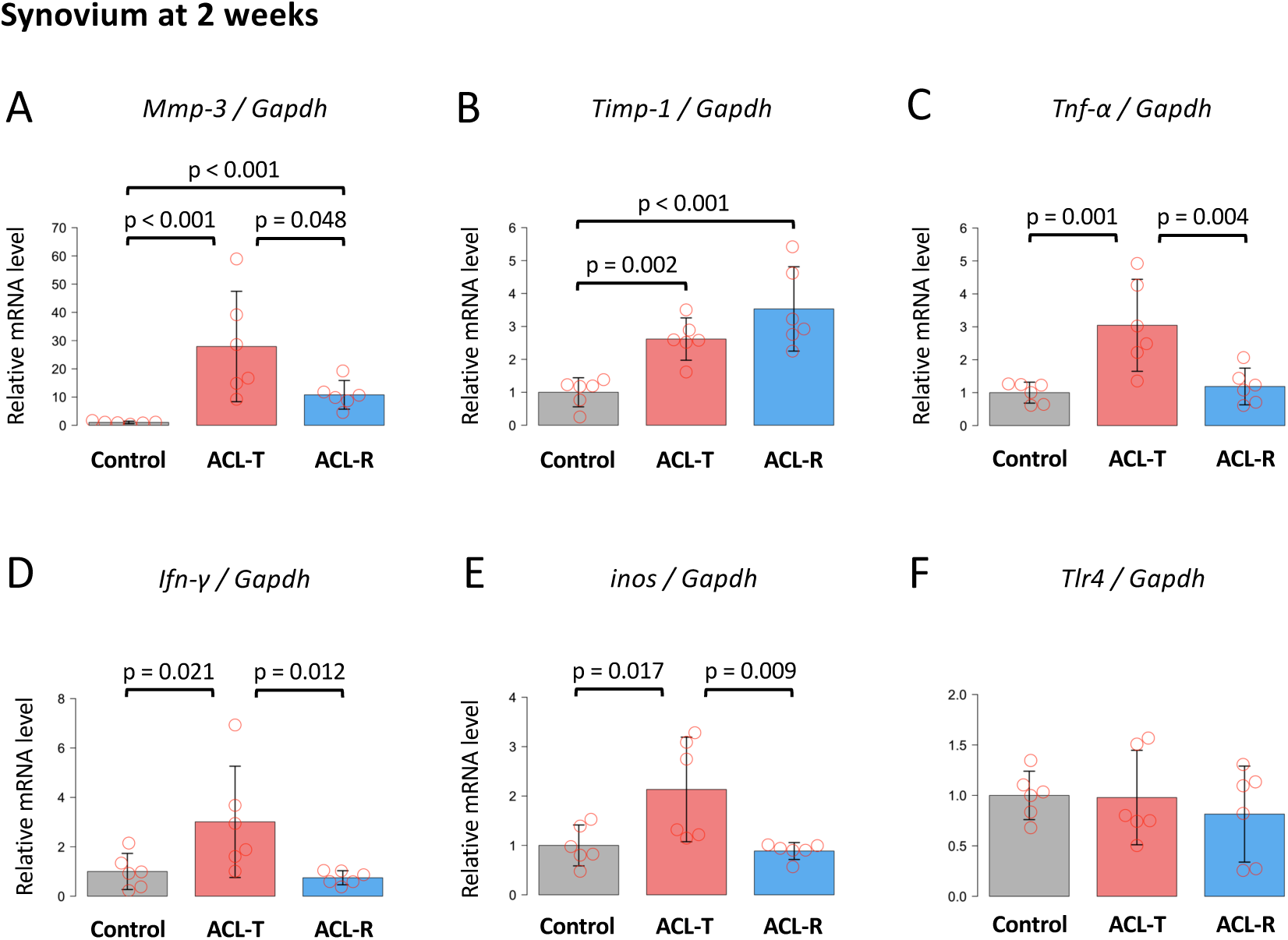
(A) Real time PCR analysis for synovium at 2 weeks. (A) *Mmp-3* expression in the ACL-T and ACL-R groups were significantly higher than that in the Control group. Moreover, the ACL-T group also significantly increased it compared to the ACL-T group. (B) *Timp-1* in both ACL injury groups was upregulated compared to that in the Control group. (C-E) The ACL-T group exhibited a significant increase of *Tnf-α, Ifn-γ, and inos* compared to the Control and ACL-R group. (F) No difference was observed in the *Tlr4* expression level between groups. Data are presented as the mean ± 95% CI.

In summary, the RT-qPCR results showed the increased expression of the *Mmp-3*, *Timp-1* of *Mmp-3* inhibitor ^30^, and *Tnf-α* in cartilage of the ACL-R group, although it induced less synovitis and cartilage degeneration. Unlike the results of cartilage, synovium in the ACL-T group showed upregulation of *Ifn-γ,* M1 macrophage polarization factor, and *iNos*, M1 macrophage marker^31–34^, in addition to *Mmp-3* and *Tnf-α*. However, the *Mmp-3* and *Tnf-α* expressions were not increased at 6 and 10 weeks.

### The expression of miR145-5p and miR149-5p in the synovial fluid increased in the ACL-T group

To verify the effect of using biomaterial for dPCR, we initially analyzed 5S rRNA expression in only Tetra-Slime as a negative control. Compared with the Tetra-slime including synovial fluid, concentration of 5s rRNA in only Tetra-Slime group was obviously lower although Unisp6 of spike-in control was observed (Supplementary Figure 5). Next, we evaluated the efficiency of collecting synovial fluid and the effect of acidic pH on miRNA expression. Digital PCR revealed that UniSp2 expression was lower in the synovial fluid group than in the Control group, and in the pH 3.0 solution, it showed reduced expression compared to the pH 7.0 solution, respectively (Supplementary Figure 6). The expression of miR145-5p^35^ and miR149-5p^36, 37^, which are downregulated by TNF-α, was investigated in the mouse synovial fluid. dPCR results showed the increased expression of *miR145-5p* and *miR149-5p* in the ACL-T group compared with other groups at 2 weeks, although this difference was not statistically significant (Fig. 7B, C).

**Fig. 7.**
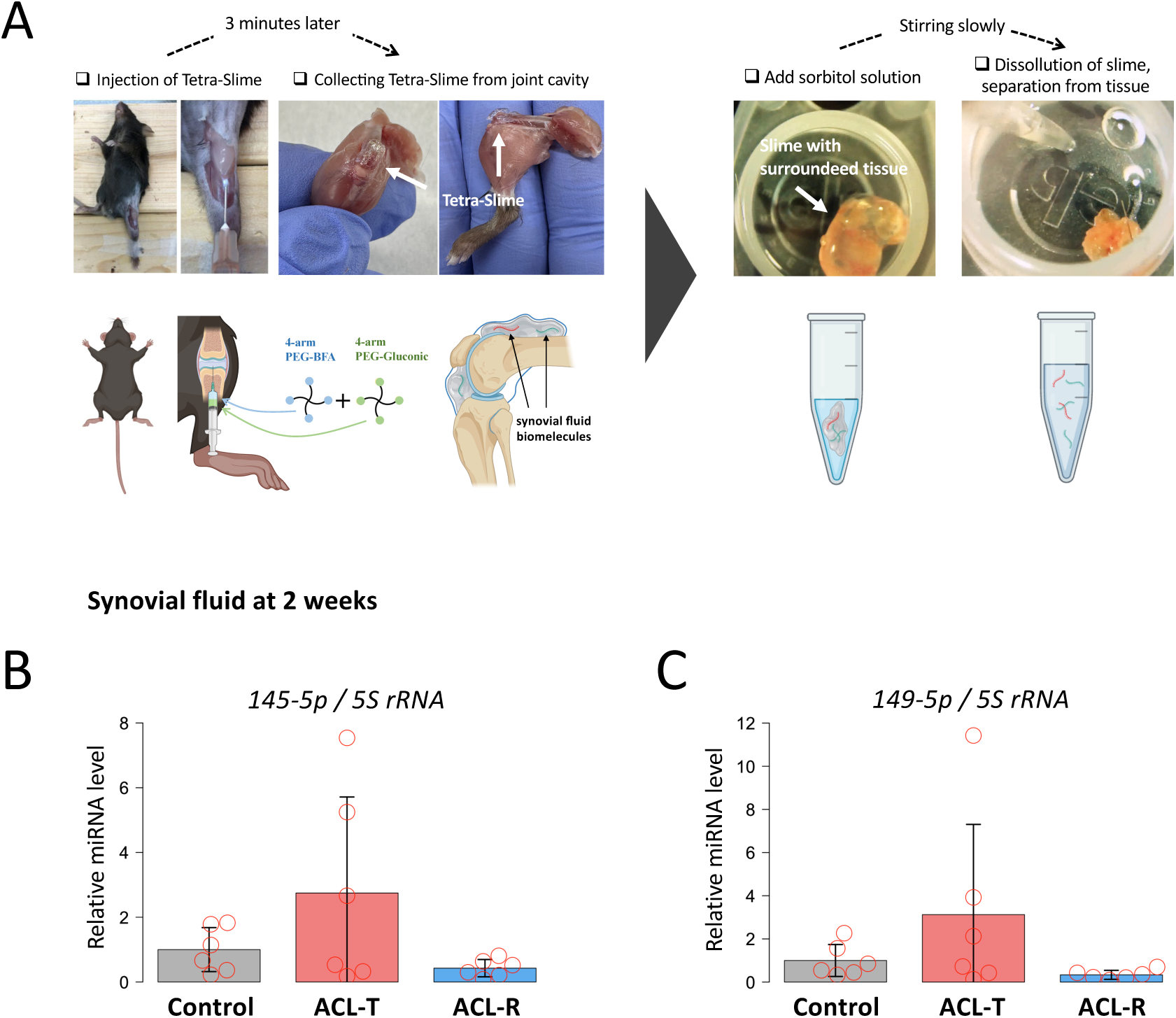
(A) Conceptual diagram of a novel method to collect synovial fluid biomolecules in the murine knee joints using Tetra-Slime. After dissolving 4-arm PEG-BFA and 4-arm PEG-Gluconic in 50 mM phosphate-buffered saline (pH 3.0) and adjusting the concentration to 0.2 g/ml, 50 µl was injected into the knee joint. 3 minutes later, solidified Tetra-Slime including synovial fluid was collected and it was resolved by 100 µL of sorbitol solution with a 0.5 g/ml concentration to separate from the surrounding intra-articular tissue. (B-C) The ACL-T group exhibited an increase of miR145-5p and miR149-5p expression in synovial fluid compared to the Control and ACL-R groups at 2 weeks. Data are presented as the mean ± 95% CI.

## Discussion

Acute synovitis and chronic mechanical stress influence PTOA progression; however, the impact of these two factors on cartilage degeneration and their underlying mechanisms remain unclear. In this study, we aimed to investigate each effect on PTOA using Invasive and Non-Invasive ACL injury mouse models. Furthermore, to elucidate tissue interaction between acute synovitis and secondary cartilage degeneration, we proposed a novel approach for synovial fluid analysis in mouse knee joint using Tetra-Slime. Our data clarified that acute synovitis in the ACL-T model induced secondary cartilage degeneration in the early stage compared to the ACL-R model. During this mechanism, real-time PCR revealed that *Mmp-3*, *Tnf-α*, and M1 macrophage activation in synovium might be involved.

We successfully established a novel method for analyzing murine synovial fluid and suggested that circulating miR145-5p and miR149-5p were possibly secreted into synovial fluid to prevent secondary cartilage degeneration induced by acute synovitis. Unlike the early stage, histological changes about synovitis and cartilage degeneration were identical between both models in the late stage, suggesting that continuous mechanical stress was the critical factor in the late stage.

Mechanical stress plays a significant role in maintaining intra-articular homeostasis; however, excessive stress induces synovitis and cartilage degeneration. During the early stage of OA progression, synovitis is generally initiated and driven by damage-associated molecular patterns (DAMPs) released from the extracellular matrix to the synovial fluid.^38–41^ DAMPs activate TLRs in the synovium, leading to cartilage degeneration through the inflammatory mediators and MMPs.^42, 43^ In this study, we demonstrated that the ACL-T group significantly induced synovitis at 2 weeks compared to the ACL-R group although there was no difference in joint instability between groups. Furthermore, real-time PCR revealed no differences in *Tlr4* expression, the receptor for DAMPs ^43^ in the synovium, among all groups. Considering that synovitis is generally induced by DAMPs through cartilage degeneration, our result indicates that acute synovitis at 2 weeks in the ACL-T group is attributed to the surgical intervention, not secondary pathogenesis. Previous studies reported that cartilage degeneration following ACL injury initiates in the medial compartment due to joint instability. ^44–46^ However, the OARSI score in the lateral compartment of the ACL-T group was significantly higher than that of the ACL-R group at 2 weeks, suggesting that acute synovitis might cause secondary cartilage degeneration in the early stage.

We revealed the effect of acute synovitis on cartilage degeneration during the OA progression, then performed molecular biological analysis to examine the tissue interaction between synovium and cartilage. MMP-3 in the synovium of OA patients was positively related to the severity ^47^, and it degrades cartilage ECM and activates other serine proteases besides MMP-1, MMP-9, and MMP-13. ^48^ Previous studies have reported that the depletion of synovial macrophages suppresses MMP-3 expression both in vivo ^49^ and in vitro. ^50^ Our data showed that *Mmp-3*, *Tnf-α*, *Ifn-γ*, and *inos* were upregulated in the synovium of the ACL-T group at 2 weeks, indicating that M1 macrophage may be involved in the tissue interaction between acute synovitis and secondary cartilage degeneration. However, macrophage function shifts based on the M1/M2 ratio^31,32^, and we only investigated the IL-10 which is anti-inflammation cytokine related to the M2 macrophage. Our data showed no significant differences in *Il-10* expression were observed between the ACL-T and ACL-R groups at every timepoints, suggesting that anti-inflammation cytokines might not be involved with this interaction. Further experiments to assess the M2 macrophage markers such as Arg-1 and Il-4 ^31^ would provide an accurate mechanism of secondary cartilage degeneration induced by acute synovitis in terms of macrophage activation.

Interestingly, the expression of Mmp-3 and Tnf-α in the cartilage increased significantly in the ACL-R group with less OA severity. It has been reported that MMP-3 and TNF-a are upregulated in the cartilage during the early stage of OA, and their expression level decrease in the late stage. ^51, 52^ Based on histological data showing early cartilage degeneration in the ACL-T group at 2 weeks, we assume that MMP-3 and TNF-a were upregulated in cartilage prior to 2 weeks, which led to early cartilage degeneration in the ACL-T model, and then downregulated by 2 weeks. Meanwhile, the ACL-R model showed identical histological changes, including cartilage degeneration and synovitis, as in the ACL-T model at 6 weeks, despite no cartilage damage and no molecular changes in synovium and synovial fluid at 2 weeks. These results indicate that abnormal mechanical stress initially induces articular cartilage degeneration rather than intra-articular inflammation, via surrounding tissue degeneration reactions such as synovial tissue and synovial fluid, which supports our previous study. ^26^ It is also suggested that the onset mechanism during the OA progression in the Invasive ACL-T models, which has been reported in previous studies, could be an experimental-specific phenomenon and differ from the human pathology we aim to elucidate. We assume that previous data using the Invasive animal models are likely attributable to local inflammation, especially in the synovium, which may also affect OA progression in the DMM model. To test this hypothesis, we need to perform this novel synovial fluid analysis at many time points on several knee OA models.

Synovial fluid is the most suitable biofluid for exploring the onset mechanism of OA because it directly reflects the changes in various intra-articular tissues. The importance of miRNA epigenetic regulation as a cellular communication is becoming increasingly recognized, and many studies have recently demonstrated its effectiveness in the synovial fluid of OA patients. ^53, 54^ However, no study has investigated the changes in synovial fluid miRNA in the mouse OA model due to the difficulty of collection and analysis. Here, we established a novel method to collect synovial fluid effectively from the murine knee joint using Tetra-Slime. dPCR showed a non-specific reaction of Tetra-Slime to 5s rRNA expression and the expression of UniSp2 spike-in RNA, meaning that it enabled the analysis of synovial fluid components accurately. Previous studies reported that TNF-α induced cartilage degeneration by downregulating miR145-5p ^35^ and miR149-5p ^36^, indicating that these miRNAs have a protective function. In the ACLT group with histological osteoarthritic changes, miR145-5p and miR149-5p in synovial fluid were increased at 2 weeks, although differences were not significant. This may be attributed to the fact that miR145-5p and miR149-5p were produced from chondrocytes to protect against secondary cartilage degeneration by acute synovitis. However, the dPCR results revealed variability in miRNA levels in the ACL-T group due to analytical difficulties and the low abundance of miRNAs in murine knee joints. To clarify the miRNA dynamics involved in OA progression, further experiments with a larger sample size are required.

In contrast to the histological OA pathologies observed at 2 weeks, there were no significant differences in the synovitis and OARSI score at 6 and 10 weeks between the ACL-T and ACL-R groups. This suggests that chronic mechanical stress with joint instability has a much greater impact on cartilage degeneration in the long term, rather than acute synovitis, in the progression of OA.

This study has three limitations: first, we could not exclude the intra-patellar fat pad (IFP) when synovium was collected from the knee joint. Therefore, the result of real-time PCR for synovium might reflect the biological changes of IFP to synovitis and mechanical stress. The second is that miRNA-derived cells were not clarified. Synovial fluid components reflect biological changes of various articular tissues, not only synovium and cartilage; therefore, we need to perform additional in vitro experiments to investigate the tissue interaction between synovium and cartilage in detail. The third is that synovial fluid molecules could not be fully collected, and acidic pH might affect miRNA integrity to some extent. However, a previous study reported that miRNA was not degraded under extreme pH condition, as miRNA is of short length, unlike other RNAs. ^55^ Therefore, we could not entirely reveal the whole signature of synovial miRNA; however, our data partially explained intra-articular change toward cartilage degeneration.

In conclusion, we successfully established a novel method for collecting synovial fluid in the murine knee joint using Tetra-slime. During the OA progression, acute synovitis led to secondary cartilage degeneration in the early stage, which might be attributed to the increase of MMP-3, TNF-α, and M1 macrophage activation in the synovium. Among this tissue interaction, synovial fluid might play a role in preventing cartilage degeneration by increasing miR145-5p and miR149-5p. Whereas in the middle and late stages, mechanical stress was a more critical factor in the progression of cartilage degeneration rather than synovitis (Fig. 8).

**Fig. 8.**
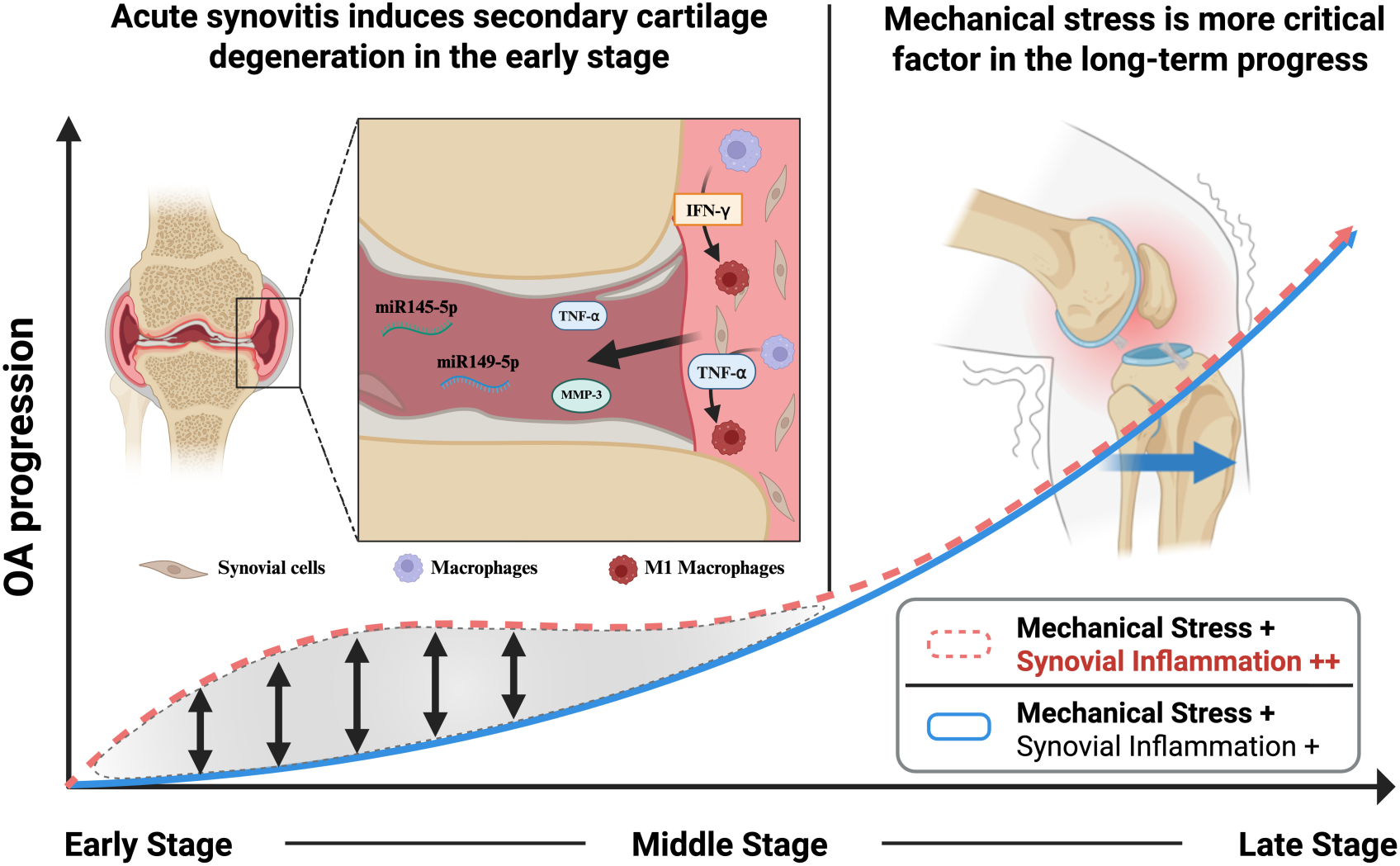
Schematic mechanism of OA development with acute synovitis and chronic mechanical stress. Acute synovitis caused secondary cartilage degeneration in the early stage and MMP-3, TNF-α, and M1 macrophage activation in the synovium were involved in this tissue interaction. Furthermore, miR145-5p and miR149-5p were secreted into the synovial fluid to protect secondary cartilage deterioration during the same time. Unlike the early stage, continuous mechanical stress had a much greater impact on cartilage degeneration rather than synovitis after the middle stage.

## Supporting information

supplemental results

## Author contributions

K.T.: data curation, investigation, methodology, validation, formal analysis, funding acquisition, writing – original draft. writing – review and editing. K.A.: investigation, methodology. T.Y., T.K, T.S.: methodology and supervision. S.E.: investigation. T.K.: project administration, resources, supervision, writing – original draft, writing – review and editing. All authors read and approved the final manuscript.

## Acknowledgments

The author(s) received no financial support for the research, authorship, and/or publication of this article. The study design and summary scheme were created with BioRender.com.

## Data Availability Statement

The data that supports the findings of this study are available in the article or from the corresponding author upon reasonable request.

## Grants or other financial supporters of the study

- Japan Society for the Promotion of Science, Research Fellows (22J23384)
- Ginff2rant-in-Aid for Japan Society for the Promotion of Science, Challenging Research (Exploratory)(21K19724)

All authors have no conflicts of interest related to the manuscript.

## References

1. Han S. Osteoarthritis year in review 2022: biology. Osteoarthritis Cartilage 2022;30:1575–82.

2. Libke ML, Cunningham DJ, Furman BD, Yi JS, Brunger JM, Kraus VB, et al. Mode of injury and level of synovitis alter inflammatory chondrocyte gene expression and associated pathways. Sci Rep 2024;14:28917.

3. Evers BJ, Van Den Bosch MHJ, Blom AB, van der Kraan PM, Koeter S, Thurlings RM. Post-traumatic knee osteoarthritis; the role of inflammation and hemarthrosis on disease progression. Front Med (Lausanne) 2022;9:973870.

4. Lieberthal J, Sambamurthy N, Scanzello CR. Inflammation in joint injury and post-traumatic osteoarthritis. Osteoarthritis Cartilage 2015;23:1825–34.

5. Peters H, Rockel JS, Little CB, Kapoor M. Synovial fluid as a complex molecular pool contributing to knee osteoarthritis. Nat Rev Rheumatol 2025;21:447–64.

6. Plsikova Matejova J, Spakova T, Harvanova D, Lacko M, Filip V, Sepitka R, et al. A Preliminary Study of Combined Detection of COMP, TIMP-1, and MMP-3 in Synovial Fluid: Potential Indicators of Osteoarthritis Progression. Cartilage 2021;13:1421S-30S.

7. Rajandran SN, Ma CA, Tan JR, Liu J, Wong SBS, Leung YY. Exploring the Association of Innate Immunity Biomarkers With MRI Features in Both Early and Late Stages Osteoarthritis. Front Med (Lausanne) 2020;7:554669.

8. Tavallaee G, Rockel JS, Lively S, Kapoor M. MicroRNAs in Synovial Pathology Associated With Osteoarthritis. Front Med (Lausanne) 2020;7:376.

9. Cortez MA, Bueso-Ramos C, Ferdin J, Lopez-Berestein G, Sood AK, Calin GA. MicroRNAs in body fluids--the mix of hormones and biomarkers. Nat Rev Clin Oncol 2011;8:467–77.

10. Li YH, Tavallaee G, Tokar T, Nakamura A, Sundararajan K, Weston A, et al. Identification of synovial fluid microRNA signature in knee osteoarthritis: differentiating early- and late-stage knee osteoarthritis. Osteoarthritis Cartilage 2016;24:1577–86.

11. Xie W, Su W, Xia H, Wang Z, Su C, Su B. Synovial Fluid MicroRNA-210 as a Potential Biomarker for Early Prediction of Osteoarthritis. Biomed Res Int 2019;2019:7165406.

12. Boffa A, Merli G, Andriolo L, Lattermann C, Salzmann GM, Filardo G. Synovial Fluid Biomarkers in Knee Osteoarthritis: A Systematic Review and Quantitative Evaluation Using BIPEDs Criteria. Cartilage 2021;13:82S–103S.

13. Karsdal MA, Christiansen C, Ladel C, Henriksen K, Kraus VB, Bay-Jensen AC. Osteoarthritis--a case for personalized health care? Osteoarthritis Cartilage 2014;22:7–16.

14. Kuyinu EL, Narayanan G, Nair LS, Laurencin CT. Animal models of osteoarthritis: classification, update, and measurement of outcomes. J Orthop Surg Res 2016;11:19.

15. Takahata K, Lin YY, Osipov B, Arakawa K, Enomoto S, Christiansen BA, et al. Concurrent joint contact in anterior cruciate ligament injury induces cartilage micro-injury and subchondral bone sclerosis, resulting in knee osteoarthritis. Osteoarthritis Cartilage 2025.

16. Arakawa K, Takahata K, Enomoto S, Oka Y, Ozone K, Nakagaki S, et al. The difference in joint instability affects the onset of cartilage degeneration or subchondral bone changes. Osteoarthritis Cartilage 2022;30:451–60.

17. Sousa-Valente J, Calvo L, Vacca V, Simeoli R, Arevalo JC, Malcangio M. Role of TrkA signalling and mast cells in the initiation of osteoarthritis pain in the monoiodoacetate model. Osteoarthritis Cartilage 2018;26:84–94.

18. Keeble J, Blades M, Pitzalis C, Castro da Rocha FA, Brain SD. The role of substance P in microvascular responses in murine joint inflammation. Br J Pharmacol 2005;144:1059–66.

19. Keeble J, Russell F, Curtis B, Starr A, Pinter E, Brain SD. Involvement of transient receptor potential vanilloid 1 in the vascular and hyperalgesic components of joint inflammation. Arthritis Rheum 2005;52:3248–56.

20. Seifer DR, Furman BD, Guilak F, Olson SA, Brooks SC, 3rd, Kraus VB. Novel synovial fluid recovery method allows for quantification of a marker of arthritis in mice. Osteoarthritis Cartilage 2008;16:1532–8.

21. Hong S, Yu T, Wang Z, Lee CH. Biomaterials for reliable wearable health monitoring: Applications in skin and eye integration. Biomaterials 2025;314:122862.

22. Wang H. Biomaterials in Medical Applications. Polymers (Basel) 2023;15.

23. Trucillo P. Biomaterials for Drug Delivery and Human Applications. Materials (Basel) 2024;17.

24. Katashima T, Kudo R, Naito M, Nagatoishi S, Miyata K, Chung UI, et al. Experimental Comparison of Bond Lifetime and Viscoelastic Relaxation in Transient Networks with Well-Controlled Structures. ACS Macro Lett 2022;11:753–9.

25. Fang G, Bian Z, Liu D, Wu G, Wang H, Wu Z, et al. Water-soluble diboronic acid-based fluorescent sensors recognizing d-sorbitol. New Journal of Chemistry 2019;43:13802–9.

26. Takahata K, Arakawa K, Enomoto S, Usami Y, Nogi K, Saitou R, et al. Joint instability causes catabolic enzyme production in chondrocytes prior to synovial cells in novel non-invasive ACL ruptured mouse model. Osteoarthritis Cartilage 2022.

27. Glasson SS, Chambers MG, Van Den Berg WB, Little CB. The OARSI histopathology initiative - recommendations for histological assessments of osteoarthritis in the mouse. Osteoarthritis Cartilage 2010;18 Suppl 3:S17–23.

28. Krenn V, Morawietz L, Burmester GR, Kinne RW, Mueller-Ladner U, Muller B, et al. Synovitis score: discrimination between chronic low-grade and high-grade synovitis. Histopathology 2006;49:358–64.

29. Livak KJ, Schmittgen TD. Analysis of relative gene expression data using real-time quantitative PCR and the 2(-Delta Delta C(T)) Method. Methods 2001;25:402–8.

30. Takahashi K, Goomer RS, Harwood F, Kubo T, Hirasawa Y, Amiel D. The effects of hyaluronan on matrix metalloproteinase-3 (MMP-3), interleukin-1a(IL-1a), and tissue inhibitor of metalloproteinase-1 (TIMP-1) gene expression during the development of osteoarthritis. Osteoarthritis Cartilage 1999;2:182–90.

31. Zhang H, Cai D, Bai X. Macrophages regulate the progression of osteoarthritis. Osteoarthritis Cartilage 2020;28:555–61.

32. Wu CL, Harasymowicz NS, Klimak MA, Collins KH, Guilak F. The role of macrophages in osteoarthritis and cartilage repair. Osteoarthritis Cartilage 2020;28:544–54.

33. Fernandes TL, Gomoll AH, Lattermann C, Hernandez AJ, Bueno DF, Amano MT. Macrophage: A Potential Target on Cartilage Regeneration. Front Immunol 2020;11:111.

34. Zhao K, Ruan J, Nie L, Ye X, Li J. Effects of synovial macrophages in osteoarthritis. Front Immunol 2023;14:1164137.

35. Hu G, Zhao X, Wang C, Geng Y, Zhao J, Xu J, et al. MicroRNA-145 attenuates TNF-alpha-driven cartilage matrix degradation in osteoarthritis via direct suppression of MKK4. Cell Death Dis 2017;8:e3140.

36. Santini P, Politi L, Vedova PD, Scandurra R, Scotto d’Abusco A. The inflammatory circuitry of miR-149 as a pathological mechanism in osteoarthritis. Rheumatol Int 2014;34:711–6.

37. Chiu YS, Bamodu OA, Fong IH, Lee WH, Lin CC, Lu CH, et al. The JAK inhibitor Tofacitinib inhibits structural damage in osteoarthritis by modulating JAK1/TNF-alpha/IL-6 signaling through Mir-149-5p. Bone 2021;151:116024.

38. de Lange-Brokaar BJ, Ioan-Facsinay A, van Osch GJ, Zuurmond AM, Schoones J, Toes RE, et al. Synovial inflammation, immune cells and their cytokines in osteoarthritis: a review. Osteoarthritis Cartilage 2012;20:1484–99.

39. Nefla M, Holzinger D, Berenbaum F, Jacques C. The danger from within: alarmins in arthritis. Nat Rev Rheumatol 2016;12:669–83.

40. Lopes EBP, Filiberti A, Husain SA, Humphrey MB. Immune Contributions to Osteoarthritis. Curr Osteoporos Rep 2017;15:593–600.

41. Lambert C, Zappia J, Sanchez C, Florin A, Dubuc JE, Henrotin Y. The Damage-Associated Molecular Patterns (DAMPs) as Potential Targets to Treat Osteoarthritis: Perspectives From a Review of the Literature. Front Med (Lausanne) 2020;7:607186.

42. Liu-Bryan R. Synovium and the innate inflammatory network in osteoarthritis progression. Curr Rheumatol Rep 2013;15:323.

43. Bartels YL, van Lent P, van der Kraan PM, Blom AB, Bonger KM, van den Bosch MHJ. Inhibition of TLR4 signalling to dampen joint inflammation in osteoarthritis. Rheumatology (Oxford) 2024;63:608–18.

44. Pauly HM, Larson BE, Coatney GA, Button KD, DeCamp CE, Fajardo RS, et al. Assessment of cortical and trabecular bone changes in two models of post-traumatic osteoarthritis. J Orthop Res 2015;33:1835–45.

45. Maerz T, Newton MD, Kurdziel MD, Altman P, Anderson K, Matthew HW, et al. Articular cartilage degeneration following anterior cruciate ligament injury: a comparison of surgical transection and noninvasive rupture as preclinical models of post-traumatic osteoarthritis. Osteoarthritis Cartilage 2016;24:1918–27.

46. Gilbert SJ, Bonnet CS, Stadnik P, Duance VC, Mason DJ, Blain EJ. Inflammatory and degenerative phases resulting from anterior cruciate rupture in a non-invasive murine model of post-traumatic osteoarthritis. J Orthop Res 2018;36:2118–27.

47. Chen JJ, Huang JF, Du WX, Tong PJ. Expression and significance of MMP3 in synovium of knee joint at different stage in osteoarthritis patients. Asian Pac J Trop Med 2014;7:297–300.

48. Murphy G, Nagase H. Progress in matrix metalloproteinase research. Mol Aspects Med 2008;29:290–308.

49. Blom AB, van Lent PL, Libregts S, Holthuysen AE, van der Kraan PM, van Rooijen N, et al. Crucial role of macrophages in matrix metalloproteinase-mediated cartilage destruction during experimental osteoarthritis: involvement of matrix metalloproteinase 3. Arthritis Rheum 2007;56:147–57.

50. Bondeson J, Wainwright SD, Lauder S, Amos N, Hughes CE. The role of synovial macrophages and macrophage-produced cytokines in driving aggrecanases, matrix metalloproteinases, and other destructive and inflammatory responses in osteoarthritis. Arthritis Res Ther 2006;8:R187.

51. Aigner T, Zien A, Gehrsitz A, Gebhard PM, McKenna L. Anabolic and catabolic gene expression pattern analysis in normal versus osteoarthritic cartilage using complementary DNA-array technology. Arthritis Rheum 2001;44:2777–89.

52. Murata K, Kanemura N, Kokubun T, Fujino T, Morishita Y, Onitsuka K, et al. Controlling joint instability delays the degeneration of articular cartilage in a rat model. Osteoarthritis Cartilage 2017;25:297–308.

53. Munjal A, Bapat S, Hubbard D, Hunter M, Kolhe R, Fulzele S. Advances in Molecular biomarker for early diagnosis of Osteoarthritis. Biomol Concepts 2019;10:111–9.

54. Palma AD, Cheleschi S, Pascarelli NA, Tenti S, Galeazzim M, Fioravanti A. Do microRNAs have a key epigenetic role in osteoarthritis and in mechanotransduction? Clin Exp Rheumatol 2016;35 518–26.

