## supplemental results for "Novel biomaterial-based synovial fluid analysis reveals protective microRNA signatures in a mouse model of acute synovitis-driven osteoarthritis"

**ACL-T and ACL-R groups increased similar joint instability**

The anterior displacement in the ACL-T and ACL-R groups was significantly higher than that in the Control groups at 2 weeks (ACL-T vs. Control, p < 0.001; CI = [1.009, 1.397]; ACL-R vs. Control, p < 0.001; CI = [1.116, 1.503]), 6 weeks (ACL-T vs. Control, p < 0.001; CI = [0.944, 1.231]; ACL-R vs. Control, p < 0.001; CI = [ 0.948, 1.235]), and 10 weeks (ACL-T vs. Control, p < 0.001; CI = [0.860, 1.159]; ACL-R vs. Control, p < 0.001; CI = [0.92, 1.219]).

**ACL-R group induced catabolic factors and inflammation cytokine expression in the articular cartilage**

In the medial tibial cartilage compared with the Control group, ACL-T group significantly increased the expression of *Mmp-3* (p < 0.001, CI = [-4.352, -2.164]), *Timp-1* (p = 0.009, CI = [ -3.714, -0.517]), and *Tnf-α* (p = 0.008, CI = [-3.271, -0.472]). ACL-R group also significantly exhibited an increase of *Mmp-3* (p < 0.001, CI = [-5.351, -3.163]), *Timp-1* (p < 0.001, CI = [-4.419, -1.222]), and *Tnf-α* (p < 0.001, CI = [-5.287, -2.488]) compared with the Control group. Moreover, *Tnf-α* in the ACL-R group was significantly higher than ACL-T group (p = 0.005, CI = [-3.415, -0.616]). In the lateral tibial cartilage, ACL-T group significantly exhibited an increase of *Mmp-3* (p < 0.001, CI = [-5.534, -2.48]) and *Timp-1* (p = 0.036, CI = [-2.934, -0.087]) compared with the Control group. *Mmp-3*, *Timp-1*, and *Tnf-α* in the ACL-R group significantly were upregulated compared with the Control group (*Mmp-3*; p < 0.001, CI = [-7.213, -4.159], *Timp-1*; p < 0.001, CI = [-4.447, -1.6], *Tnf-α*; p = 0.008, CI = [-4.245, -0.633]) and ACL-T group (*Mmp-3*; p = 0.03, CI = [-3.206, -0.151], *Timp-1*; p = 0.036, CI = [-2.936, -0.089], *Tnf-α*; p < 0.001, CI = [-5.167, -1.555]).

*Tgf-β1* decreased significantly in the ACL-R group compared with the Control (p < 0.001, 95% CI = [0.408 to1.45]) and ACL-T groups (p = 0.002, 95% CI = [0.306 to 1.348]) (Fig. S1A). *Il-1β* and *Il-10* expression increased significantly in the ACL-T group compared with the Control group (*Il-1β*: p = 0.022, 95% CI = [-5.063 to -0.37], *Il-10*: p < 0.001, 95% CI = [-5.255 to -1.461]) (Fig. S1B, D). No significant differences were observed in the *Il-6* between groups (Fig. S1C). Whereas, no significant differences were observed in the *Tgf-β1, Il-1β, Il-6,* and *Il-10* between groups (Fig. S1E-H).

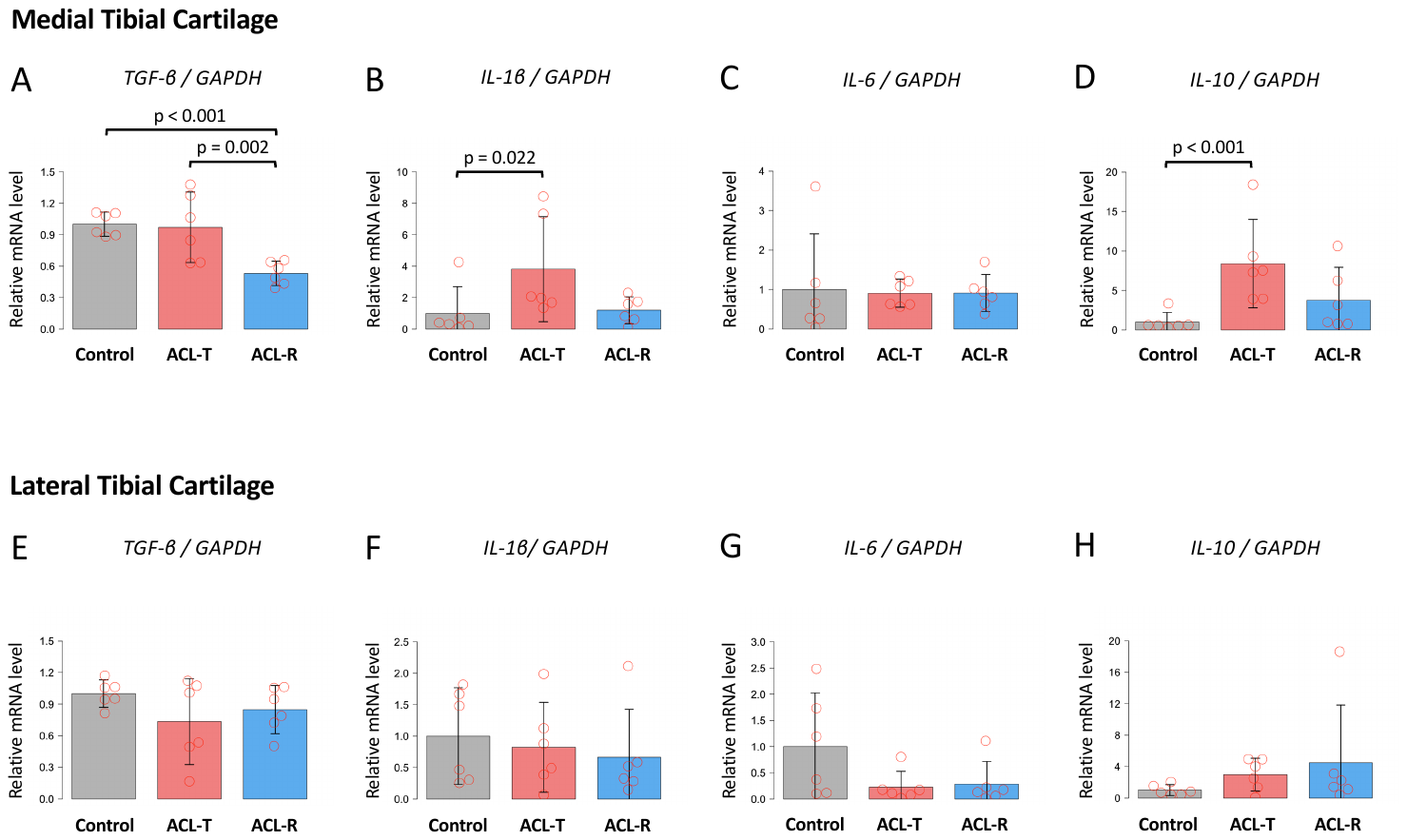

Supplementary Figure 1. Real time PCR analysis for medial and lateral tibial articular cartilage at 2 weeks. (A-D) In the medial cartilage, *Tgf-β1* expression in the ACL-R group was significantly decreased compared to that in the Control and ACL-T groups. In addition, the ACL-T group exhibited a significant increase of *Il-1β and Il-10* compared to Control group. (E-H) In the lateral cartilage, there were no significant differences in the expression between groups. Data are presented as the mean ± 95% CI.

**ACL-T group increased the expression of catabolic factors, inflammation cytokine and M1 macrophage marker in the synovium**

At 2 weeks, *Mmp-3* expression was significantly higher in the ACL-T group (p < 0.001, CI = [-4.352, -2.164]) and ACL-R group (p < 0.001, CI = [-5.351, -3.163]) than the Control group. Furthermore, ACL-T group also exhibited an increase of *Mmp-3* significantly compared with the ACL-R group (p = 0.048, CI = [0.007, 2.437]). *Timp-1* was elevated significantly in the ACL-T group (p = 0.002, CI = [-2.500, -0.576]) and ACL-R group (p < 0.001, CI = [-2.901, -0.977]) compared with the Control group. ACL-T group significantly increased *Tnf-α*, *Ifn-γ*, and *inos* compared with the Control (*Tnf-α*; p = 0.001, CI = [-2.461, -0.629], *Ifn-γ*; p = 0.021, CI = [-3.097, -0.236], *inos*; p = 0.017, CI = [-1.911, -0.186]) and ACL-R group (*Tnf-α*; p = 0.004, CI = [0.439, 2.27], *Ifn-γ*; p = 0.012, CI = [0.389, 3.249], *inos*; p = 0.009, CI = [-2.008, -0.282]). However, no significant difference was observed in the *Tlr4* expression between all groups.

No significant differences were observed in the *Tgf-β1* between groups (Fig. S2A). ACL-T group significantly increased *Il-1β* compared with the Control group (p = 0.002, 95% CI = [-7.615 to -1.697]) (Fig. S2B). *Il-6* increased significantly in the ACL-T (p < 0.001, 95% CI = [-6.833 to -2.651]) and ACL-R groups (p = 0.006; 95% CI = [-5.004 to -0.823]) compared with the Control group (Fig. S2C). Moreover, *Il-10* exhibited a significant increase in the ACL-T (p = 0.035, 95% CI = [-5.244 to -0.177]) and ACL-R groups (p = 0.026, 95% CI = [-5.396 to -0.330]) compared with the Control group (Fig. S2D).

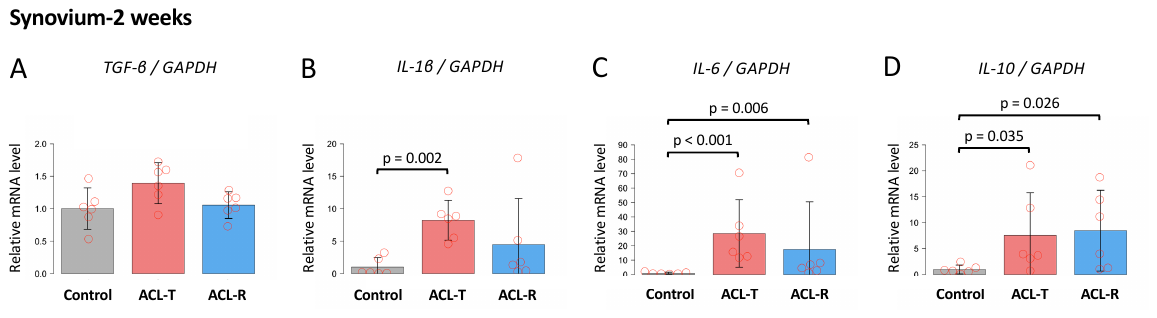

Supplementary Figure 2. Real time PCR analysis for synovium at 2 weeks. (A-D) *Il-1β, Il-6, and Il-10* expression in the ACL-T group was significantly increased compared to that in the Control group. Furthermore, the ACL-R group exhibited a significant increase of *Il-6 and Il-10* compared to Control group. Data are presented as the mean ± 95% CI.

At 6 weeks, *Mmp-3* expression significantly increased in the ACL-T (p < 0.001, 95% CI = [-4.047 to -1.676]) and ACL-R groups (p = 0.003, 95% CI = [-2.994 to -0.623]) (Fig S3A). *Timp-1* also significantly increased in the ACL-T (p = 0.049, 95% CI = [-2.544 to -0.004]) and ACL-R groups (p = 0.001; 95% CI = [-3.334 to -0.796]) (Fig S3B). ACL-T group significantly increased *Ifn-γ* compared with the Control (p = 0.02, 95% CI = [-2.705 to -0.219]) and ACL-R groups (p < 0.001, 95% CI = [0.984 to 3.469]) (Fig S3H). However, no significant differences were observed in the expression of *Tgf-β1*, *Tnf-a*, *Il-1β*, *Il-6*, and *Il-10*.

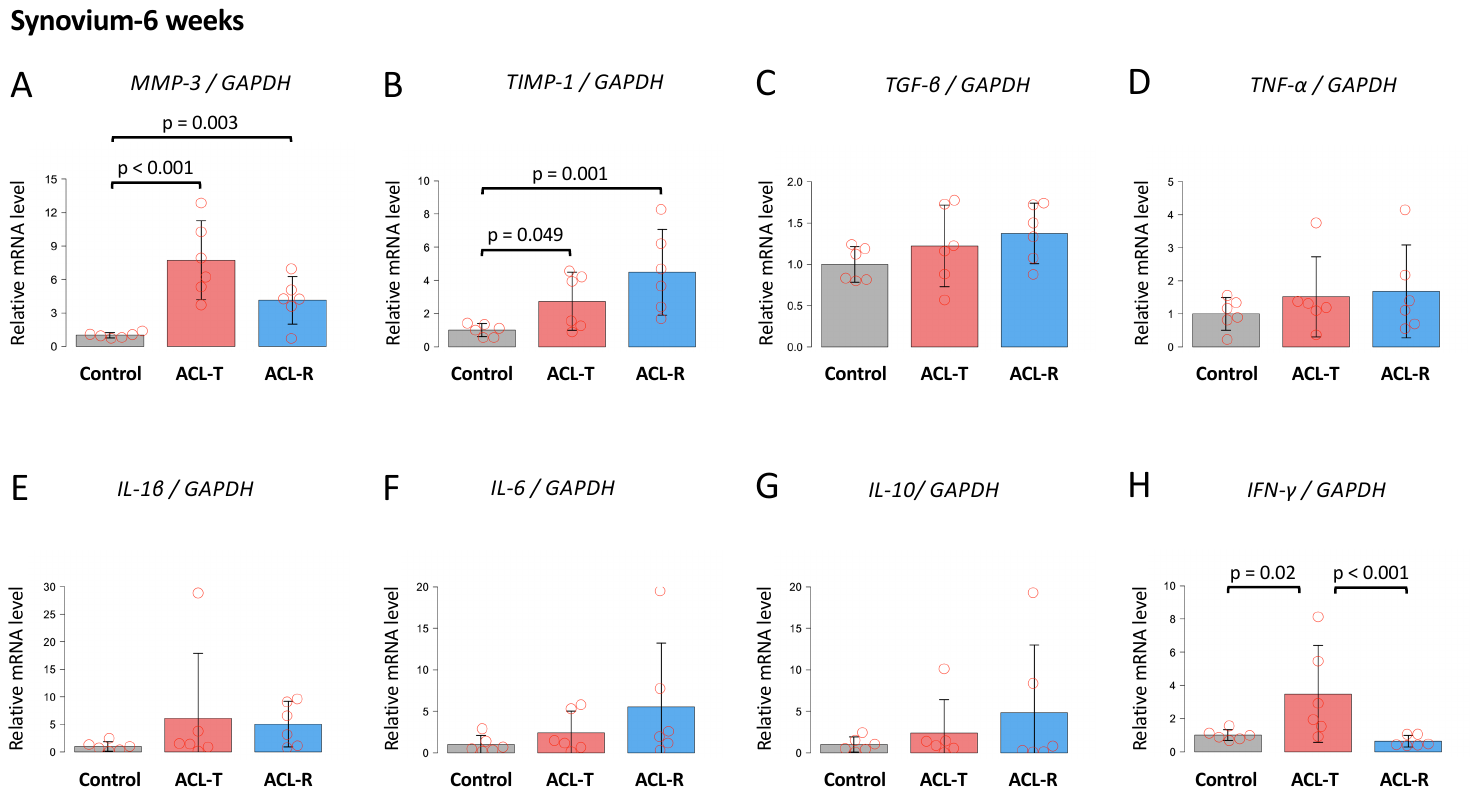

Supplementary Figure 3. Real time PCR analysis for synovium at 6 weeks. (A-B) *Mmp-3* and *Timp-1* in the ACL-T and ACL-R groups was significantly higher than that in the Control group. Moreover, the ACL-R group exhibited a significant increase of *Ifn-γ* compared to Control and ACL-R groups. Data are presented as the mean ± 95% CI.

At 10 weeks, *Mmp-3* expression significantly increased in the ACL-T (p = 0.002, 95% CI = [-8.344 to -1.81]) and ACL-R groups (p = 0.034, 95% CI = [-6.775 to -0.241]) (Fig S4A). There were no significant differences in *Timp-1, Tgf-β1*, *Tnf-a*, *Il-1β*, *Il-6*, *Il-10*, and *Ifn-γ.*
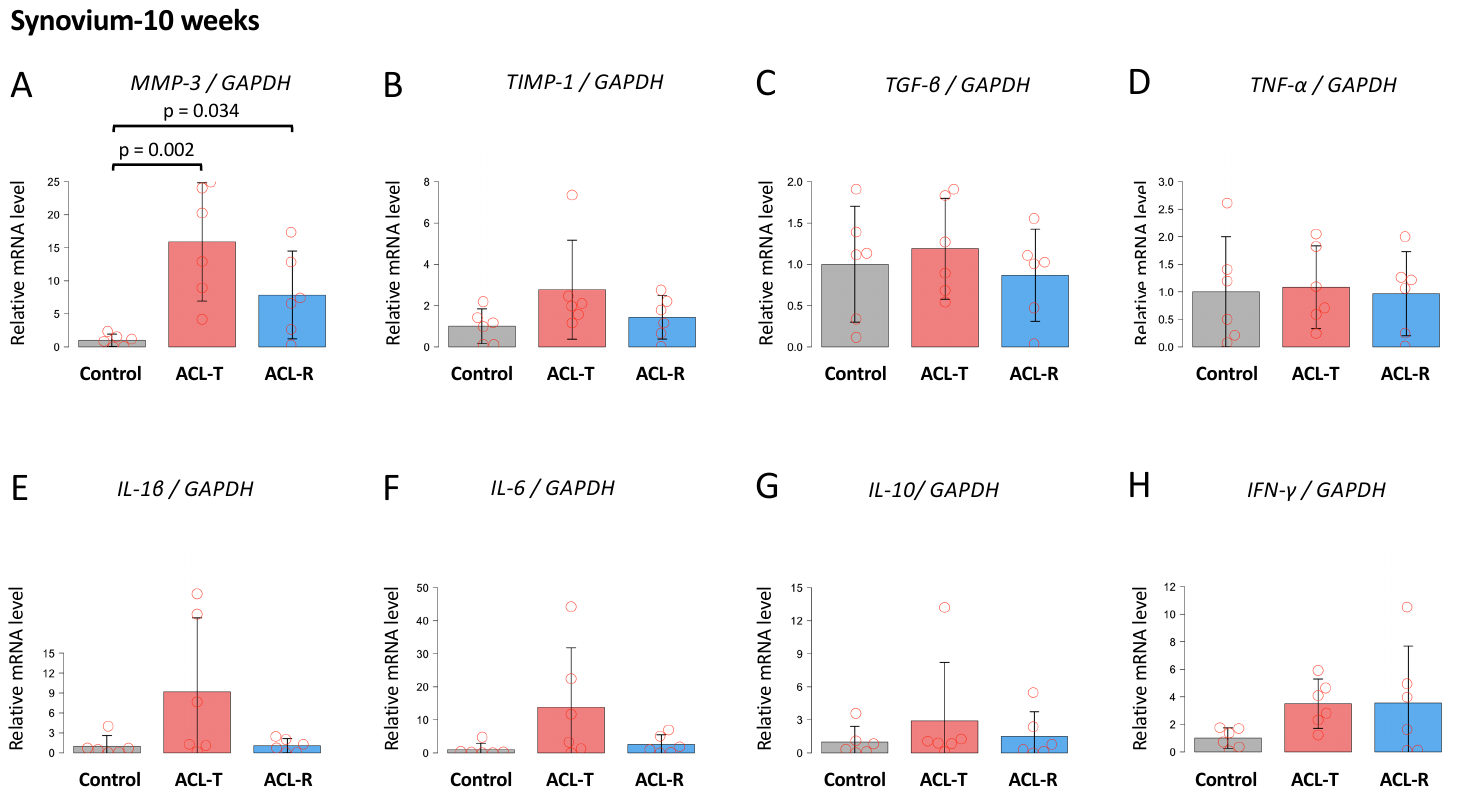

Supplementary Figure 4. Real time PCR analysis for synovium at 10 weeks. (A) *Mmp-3* in the ACL-T and ACL-R groups was significantly higher than that in the Control group. Data are presented as the mean ± 95% CI.

**No side-effect of using Tetra-Slime to collect synovial fluid in murine knee joints**

To verify the effect of using biomaterial for dPCR, we analyzed 5S rRNA expression in only Tetra-Slime as a negative control. Compared with the Tetra-slime including synovial fluid, concentration of 5s rRNA in only Tetra-Slime group was obviously lower although Unisp6 of spike-in control was observed. We also evaluated the efficiency of collecting synovial fluid and effect of acidic pH on miRNA expression. Digital PCR showed that the lower expression of UniSp2 in the synovial fluid group and pH 3.0 solution compared to the Control group and pH 7.0 solution, respectively (Fig S5-6).

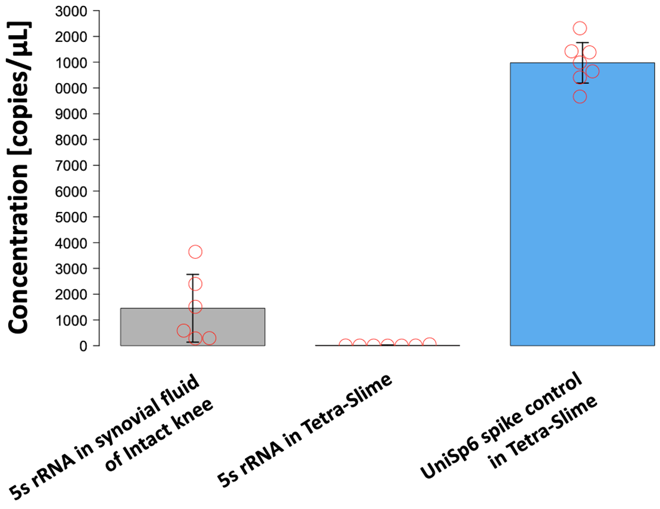

Supplementary Figure 5. Digital PCR analysis for Tetra-Slime and that including murine synovial fluid. The concentration of 5s rRNA in Tetra-Slime used to collect synovial fluid in murine knee joints was obviously higher than that in the only Tetra-Slime although Unisp6 of spike-in control was observed. Data are presented as the mean ± 95% CI.

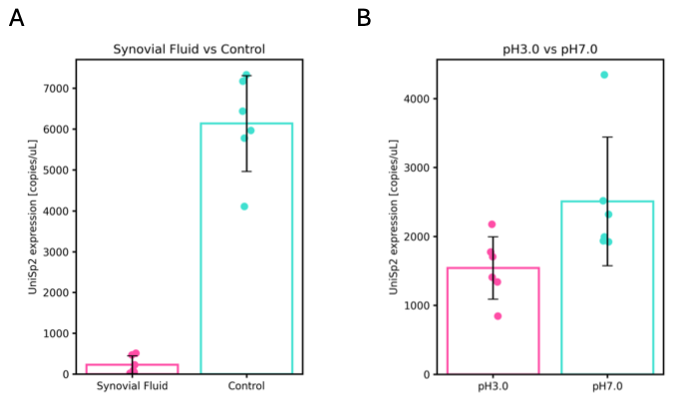

Supplementary Figure 6. (A). Digital PCR analysis for spike-in RNA in the synovial fluid group using the Tetra-Slime vs. control group. The expression of UniSp2 was lower than that in the Control group. (B). Digital PCR analysis for spike-in RNA in the pH 3.0 vs. pH 7.0 group. pH 3.0 group exhibited a decrease of UniSp2 level compared to the pH 7.0 group. Data are presented as the mean ± 95% CI.
